# Ancient atmospheric changes could favour C_3_-to-CAM transition: A simulation-based evidence

**DOI:** 10.1101/2025.09.11.675551

**Authors:** Devlina Sarkar, Sudip Kundu

## Abstract

Crassulacean Acid Metabolism (CAM) plants have very low water demand and can survive in arid environment. It is hypothesized to evolve from C_3_ metabolism about 20 to 30 million years ago (MYA) when the Earth faced declining atmospheric CO_2_ concentration ([CO_2_]_a_), increasing aridity and decreasing temperature. Understanding whether and how these atmospheric changes favour C_3_-to-CAM transition will help in bioengineering CAM into crop plants to tackle the threat of global climate change and ensure food security. Our simulations of plant cellular metabolism under the changing atmospheric conditions of ancient times observe C_3_-to-CAM transition, capture possible temporal alterations of active metabolic pathways and further confirm that both the reduced [CO_2_]_a_ and increased water scarcity associated with higher aridity, can act as evolutionary agents, driving the C_3_-to-CAM transition. Although the predicted elevated [CO_2_]_a_ of future reveals a reversion towards C_3_-like behaviour, drought always favours CAM regardless of [CO_2_]_a_ and temperature levels. Moreover, a minimum oxygen concentration is required to sustain elevated nocturnal respiration necessary for CAM, reflected by the increased activities of enzymes involved in the TCA cycle and the mitochondrial electron transport chain.

## 1. Introduction

In C_3_ plants, CO_2_ primarily enters the leaf during the daytime and is directly fixed through the Calvin-Benson-Bassham (CBB) cycle. Whereas, in Crassulacean acid metabolism (CAM), the process of CO_2_ uptake and fixation in CBB cycle are temporally separated. The atmospheric CO_2_ enters the internal air spaces of the leaf at night, fixed into malate, a C_4_ compound, which gets stored at night and then again decarboxylated at day to provide CO_2_ to CBB cycle. This is an alternative photosynthetic pathway with low water demand due to less transpirational water loss and high adaptive capacity in extreme hot and arid regions. Engineering CAM into crop plants is a promising strategy for improving water-use efficiency (WUE) (Lim *et al*. 2025). Designing such drought-resistant crops with improved WUE is therefore crucial for ensuring food security under the predicted extreme environmental conditions in the near future (Heyduk 2022). Time-calibrated phylogenies and large-scale carbon isotope (δ^13^C) studies suggest that CAM evolved from C_3_ approximately 20-30 million years ago (MYA) (Heyduk 2022; Sage et al., 2023). Further evidences suggest that during the period preceding this evolution, spanning from early-Eocene (56 MYA) to early-Miocene (20 MYA), atmospheric CO_2_ concentration ([CO_2_]_a_) declined from ∼700-1000 ppm to ∼200 ppm (Bellasio et al., 2021; Sage et al. 2023), Earth’s surface temperature (T) decreased from ∼35ºC to ∼22ºC (Passchier *et al*. 2013; Judd *et al*. 2024); and aridification increased *(DeCelles et al. 2007; Passchier et al. 2013; Zhang et al., 2015; Wang et al. 2016)* (i.e. relative humidity (RH) decreased from an initial value of ∼67% in middle-Miocene (Jahren, Silveira & Sternberg 2003)). These atmospheric and climatic shifts are hypothesized to have favoured the evolution of CAM from C_3_ photosynthesis during that ancient time (Sage *et al*. 2023). Understanding whether and how these atmospheric changes favoured the metabolic shift from C_3_ to CAM may help the biotechnologists to engineer water-efficient crops.

Unlike C_3_, CAM plants exhibit an inverted diel pattern of stomatal opening and closing, with CO_2_ uptake occurring partially or predominantly at night. This temporal separation of gas exchange and hence stomatal opening from transpiration during the hotter daytime, substantially reduces the water loss and enhances WUE in water-limited environments. The gaseous exchange in plants occurs through microscopic pores, called stomata, which are controlled by surrounding guard cells (GCs). Stomata open when the concentration of osmolytes like K_2_malate, KCl, sucrose etc., within GC increases, leading to water influx from the surrounding subsidiary cells and a consequent rise in turgor pressure (Lawson 2009). Upon opening, CO_2_ enters into the internal air spaces of the leaf and is primarily fixed by the mesophyll cells (MCs). which, in return, supply sucrose to GC. This tightly coordinated GC-MC metabolism enables the CO_2_ uptake, its fixation into organic compound and production of phloem sap. As mentioned earlier, in CAM, CO_2_ enters the internal air spaces of the leaf at night. It is initially assimilated into oxaloacetate (OAA) by the carboxylating enzyme phosphoenolpyruvate carboxylase (PEPC), and subsequently reduced to malate by cytosolic malate dehydrogenase (MDH). This malate is stored overnight in the vacuole and is decarboxylated during the following day by major decarboxylating enzymes such as phosphoenolpyruvate carboxykinase (PEPCK) and NADP-dependent malic enzyme (NADP-ME) (Winter & Smith 2022).

Previous experimental efforts explored the physiological (Guan et al. 2020) and molecular changes (Kong *et al*. 2020), altered gene expressions (Heyduk *et al*. 2019; Shen *et al*. 2022), transcriptomic (Bräutigam et al., 2016; Ossa et al. 2022) and proteomic (Guan *et al*. 2021) changes, the roles of different phytohormones in controlling stomatal behaviour (Wakamatsu *et al*. 2021) etc., during C_3_-to-CAM transition in different facultative CAM species (Winter & Holtum 2024) under different stress conditions including salt stress, drought stress, higher irradiance etc (Tan & Chen 2023). The different expression levels of carbonic anhydrase genes in *Arabidopsis thaliana* leaves were also reported under different atmospheric CO_2_ concentrations (Rudenko et al., 2022). On the other hand, several computational efforts simulated the C_3_-to-CAM transition only in MC either by forcefully restricting the daytime CO_2_ uptake by the system (Tay et al. 2021) or gradually reducing transpirational water loss accompanied by a trade-off in phloem sap production (Töpfer et al. 2020). Recently, we developed a dual-cell (GC and MC) diel metabolic model, coupled with a gas diffusion model which links the CO_2_ uptake to transpirational water loss, depending on the temporal fluctuations of T and RH (Sarkar & Kundu 2025). Simulation showed gradual reduction of the transpirational water loss shifts the metabolism from C_3_ to CAM and at the same time, captured the temporally differential activities of several enzymes involved in different pathways of central carbon metabolism, during this transition. However, none of these studies addressed whether and how atmospheric changes during early-Eocene to early-Miocene might have driven the shift from C_3_ metabolism to CAM. To investigate this, we simulate our previously published six-phase GC-MC diel metabolic model (Sarkar & Kundu 2025) of a plant leaf using a constraint-based modelling approach (Flux Balance Analysis or FBA) (Orth et al. 2010), under varying atmospheric conditions of that geological time period with a dual objective of optimizing the phloem sap production and cellular economy. Results capture the C_3_-to-CAM transition and confirm that both the reduced [CO_2_]_a_ and increased water scarcity associated with higher aridity, can act as evolutionary agents, driving the C_3_-to-CAM transition. Additionally, the study provides a quantitative assessment of changes in flux through key reactions of central carbon metabolism during the transition, arising from either reduced [CO_2_]_a_ or enhanced aridity. While declining [CO_2_]_a_ results in beginning of nocturnal CO_2_ uptake, accompanied by increased daytime starch accumulation, enhanced nocturnal malate storage, and elevated activities of C4-like enzymes such as PEPC, PEPCK, ME, and MDH etc.; aridity not only induces all of these changes but also leads to the emergence of an inverted stomatal rhythm. Moreover, a minimum oxygen concentration is needed to support higher nocturnal respiration, necessary for CAM. While the simulation under predicted elevated [CO_2_]_a_ of the future reveals a reversion towards C_3_-like behaviour, drought always favours CAM.

## 2. Materials and methods

Here, our previously constructed, time-resolved, atmospheric factors coupled, dual-cell (GC-MC) diel metabolic model of a plant leaf (Sarkar & Kundu 2025) is used. The 24-hour diel cycle is divided into six phases to capture the change in the pattern of CO_2_ uptake and metabolite storage and production of different metabolites in different phases. To capture the CO_2_ uptake pattern at daytime in CAM (Heyduk 2022), we divided the 12-hour daytime into three phases (phases 1-3). Moreover, it has also been experimentally reported that in C_3_ plants, GC produces starch from middle phase of the day to the first phase of night (Horrer *et al*. 2016). To capture these dynamics, we further divided the 12-hour nighttime (phase IV of the CAM cycle, reported in Heyduk, 2022 (Heyduk 2022)) into three phases (phases 4-6), allowing the model to reflect the temporal metabolic and CO_2_ uptake patterns accurately. The six phases, denoted as phase 1 to 6, represent different time intervals of a diel cycle: phase 1 (0-1 h), phase 2 (1-11 h), phase 3 (11-12 h), phase 4 (12-13 h), phase 5 (13-23 h) and phase 6 (23-24 h) respectively. Thus, the 24-hour time span of a complete diel cycle is divided into these six phases in a ratio of 1:10:1:1:10:1 This six-phase dual-cell (GC-MC) metabolic model is further coupled with a gas diffusion model [see **Figure 1a**, equation (1) and equation (2)]. The transpirational water loss from the system in each phase depends on different environmental parameters such as T and RH of that particular phase. The equations of a gas diffusion model are given below.

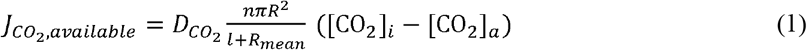

**Figure 1.**
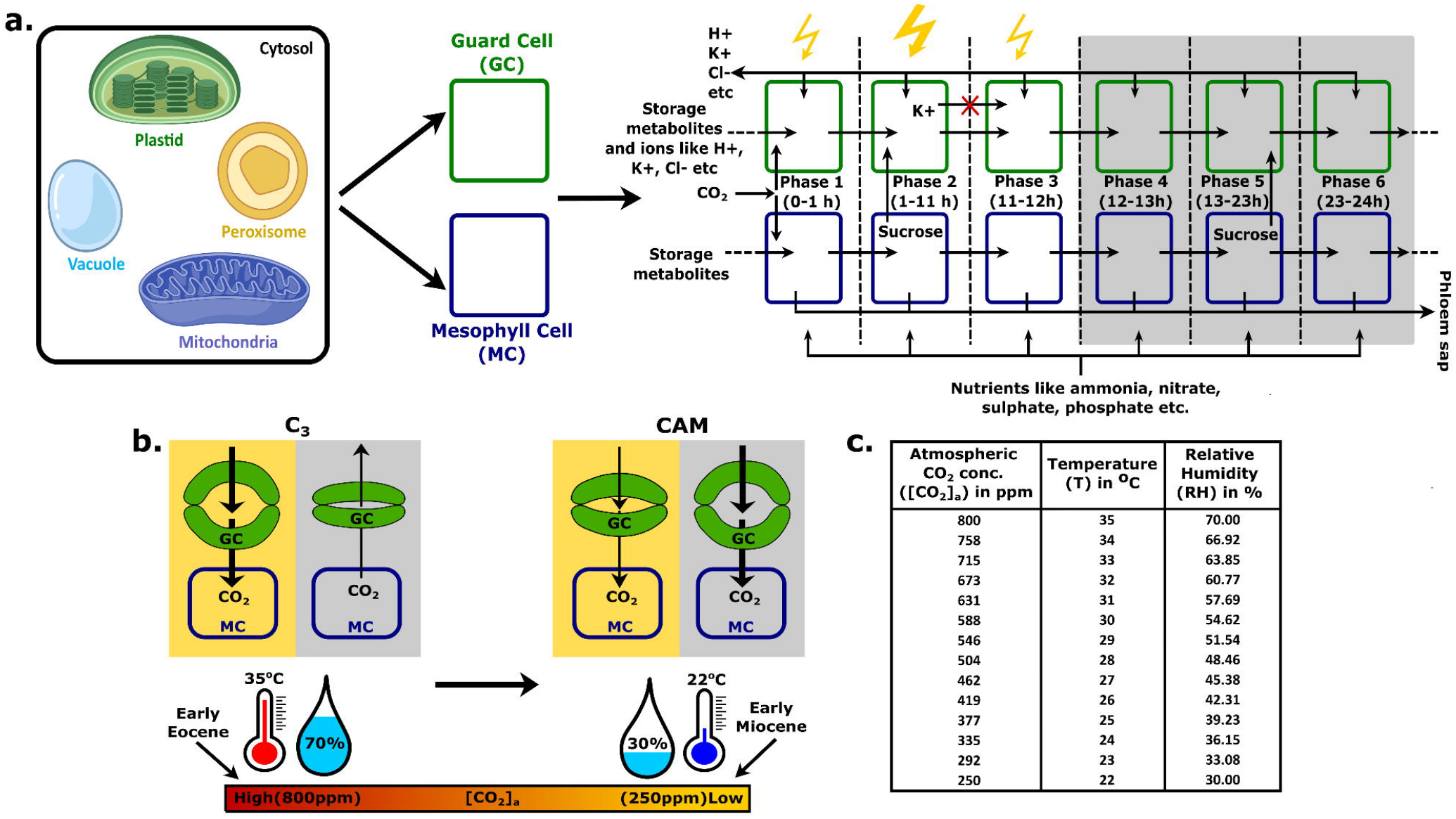
Schematic representation of the model and constraints for simulating the model. **a**. Schematic representation of the construction of the six-phase diel metabolic model comprising guard and mesophyll cells, with each cell containing different cellular compartments-cytosol, mitochondria, chloroplast, peroxisome, vacuole and endoplasmic reticulum (not shown here). Only MC can produce phloem sap. Ions can enter and exit GC in any of the six phases. Metabolites and ions are transported from one phase to another phase in such a way that one metabolite produced in one phase may get utilized in the next phase. In GC, the transfer of K^+^ from phase 2 to 3 is blocked. Images of organelles are taken from biorender.com. **b**. Schematic representation of the simulation from C_3_ to CAM under the changing environment of ancient times (early-Eocene to early-Miocene). **c**. The values of temperature (T) and relative humidity (RH) at different atmospheric CO_2_ concentration ([CO_2_]_a_) corresponding to different points between early-Eocene to early-Miocene time periods are listed.

Equation 1 represents the flux of 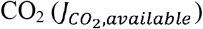 that can enter the leaf’s internal airspace through a stomatal pore of radius R at a specific concentration of atmospheric CO_2_ ([CO_2_]_a_) and T. Moreover, [CO_2_]_i_ represents the concentration of CO_2_ inside the leaf, 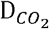 is the diffusion coefficient of CO_2_, n is the number of stomata per unit area of the leaf, πR^2^ is the average area per stomatal pore, *l* is the depth of stomatal pore and R_mean_ is the mean radius which is added to the depth of the stomatal pore as a correction factor (Nobel 2009). A plot of 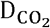 values at some particular temperatures, listed in Appendix I (Nobel 2009), gives us a straight-line graph, showed in **Supporting Information Figure S1**. We use linear regression method to get the c (intercept) and m (slope) of that straight line. Now, we can calculate 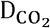 values at specific temperatures of different phases of the diel cycle in different geological time periods. In a similar manner, we plot the internal CO_2_concentration of the leaf ([CO_2_]_i_) at different atmospheric CO_2_ concentrations ([CO_2_]_a_) (>100 ppm) (Moss & Rawlins 1963) and get the c (intercept) and m (slope) values using linear regression method. Using these values, we can calculate the [CO_2_]_i_ for any specific [CO_2_]_a_ value during early-Eocene to early-Miocene time period. The radius of the stomatal pore (R) at daytime is considered to be 7 µm with a basal value of 0.6 µm in the C_3_ plant and the depth of the stomatal pore is considered to be twice the radius of the pore, assuming the pore to be circular, as used in our previously published work (Sarkar & Kundu 2025). Research suggests that species differ in how stomatal density responds to elevated 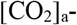 some increasing, some decreasing, and others showing none or only minor changes (Serna 2008). To simplify the simulation and focus specifically on metabolic changes driven by atmospheric conditions, initially, the number of stomata per mm^2^ is considered to be fixed at 250, which is within the reported values, ranging from 45 to 720 (Hetherington & Woodward 2003). Later, to capture the effect of varying stomatal density under changing [CO_2_]_a_, we increase/decrease the stomatal density as the atmospheric condition changes form early-Eocene to early-Miocene.

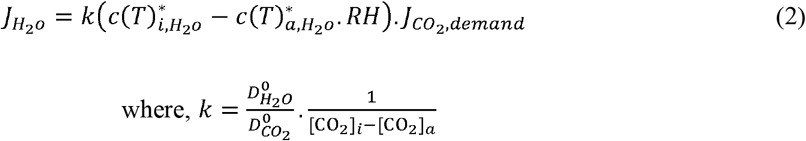

Equation 2 gives us the relationship between the water loss through transpiration and the CO_2_ demand of the system depending on the T and RH in each phase throughout the diel cycle. The values of T and RH for different phases of a diel cycle are estimated using a skewed sinus generator (Töpfer *et al*. 2020). 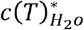 is the saturation concentration of water vapour, 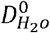 and 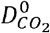 are the standard diffusion coefficients of H_2_O and CO_2_, and 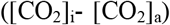 is the concentration difference of CO_2_ inside and outside the leaf. We convert the values of concentration of CO_2_ ([CO_2_]) from ppm to µmol.m^-3^ to use in our simulation. The metabolisms of GC and MC are interlinked in this model in a very simple way. The accumulation of osmolytes in GC opens stomata. Equation 1 gives us the maximum flux of CO_2_ that can enter the system depending on the size of stomatal pore, diffusion coefficients at different temperatures in different phases of the diel cycle and the concentration difference of CO_2_ inside and outside the leaf. The amount of CO_2_ entering each phase is further scaled by multiplying it with the duration of that phase. The entered CO_2_ is available to 300 MCs and 1 GC in the internal space of the leaf as the ratio of MC to GC is considered to be 300, as used in our previous study (Sarkar & Kundu 2025). On the other hand, MC supplies sucrose to GC, which ultimately helps in stomatal opening. Sucrose is allowed to transfer from MC to GC in phase 2 at daytime and in phase 5 at nighttime. The rate of solute accumulation (i.e, osmotic pressure or OP) is 1.928 µosmol.m^-2^.s^-1^ for 7 µm aperture size (Sarkar & Kundu 2025). Assuming the photosynthetic capacity of GC to be 20%, the amount of sucrose transferred to GC is 80% of the OP, as previously used (Sarkar & Kundu 2025). In the model, ions (K^+^, Cl^-^, H^+^) can move in and out of GC during any phase of the day or night. However, experimental studies have shown that K^+^ accumulates in GC primarily during early morning, while sucrose accumulation takes place later during the day (Lawson 2009). Based on this, the transfer of K^+^ from phase 2 to phase 3 is blocked in the model. The schematic representation of the model is shown in **Figure 1a**, the model files are provided in the **Supporting Information Datasets S1** and **S2**; and all the other constraints are listed in the **Supporting Information Table S1**. The constraints, which we have taken from our previously published paper (Sarkar & Kundu 2025), are based on various experimental observations (see references therein).

We have used Flux Balance Analysis (FBA) (Orth *et al*. 2010), a constraint-based modelling approach for simulating our model. In this approach, the reactions of the metabolic model are represented in the form of a stoichiometry matrix, **S** of size m x n, where m represents the number of metabolites and n represents the number of reactions present in the model. At steady state, the rate of production and the rate of consumption are equal for each internal metabolite in the model. This can be mathematically represented as,

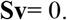

where, the vector **v** represents the flux through all the n number of reactions and **S** is the stoichiometry matrix.

The optimization problem can be defined as,

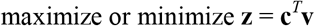

where, **z** is the objective function, **c**^*T*^ is the transpose of a vector (**c**) of weights that indicate how much a reaction contributes to the objective, **v** is the vector of all fluxes.

The constraints on the flux values are given as,

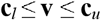

where, **v** is the vector of all fluxes, and **c**_*u*_ and **c**_*l*_ are the upper and lower bounds of fluxes, respectively.

The objective used in this study is maximization of phloem sap production, followed by the minimization of the total absolute sum of fluxes (proxy of minimization of cellular economy). The scobra package (Shaw & Cheung 2018) is used for simulations of FBA. The codes for all the simulations are given in **Supporting Information Dataset S3**.

It is reported that [CO_2_]_a_ declined from ∼700-1000 to ∼200 ppm (Bellasio *et al*. 2021; Sage *et al*. 2023), T decreased from 35-22ºC (Passchier *et al*. 2013; Judd *et al*. 2024) and RH decreased (Passchier *et al*. 2013) during the early-Eocene to early-Miocene time period. There is no report about the values of RH in early-Eocene and early-Miocene, whereas it is reported to be ∼67% in the middle Eocene (Jahren *et al*. 2003). However, several studies reported the reduction of RH during this time interval (Passchier *et al*. 2013). Therefore, we consider the value of RH in early-Eocene and early-Miocene to be 70% and 30%, slightly higher and lower than that in middle-Eocene respectively. Some known physiological parameters for C_3_ and CAM, already used in our previous work (Sarkar & Kundu 2025) are also used here, and those are listed in **Supporting Information Table S1**. Some constraints like ratio of photon values in different phases of day, flux of the photon in the middle phase (phase 2) of the day (200 µmol.m^-2^.s^-1^), percentage of sucrose transferred from MC to GC in phase 2 and phase 5 etc., remain unchanged throughout the simulation from early-Eocene to early-Miocene time period. There is no data available about the temporal variation of T and RH in ancient times and therefore, we assume the difference of max and min T and RH to be 10ºC and 10% respectively. Experimental evidences showed that the maximum stomatal apertures for basil and broad bean are 7 µm and 7.4 ± 2.3 µm respectively (Outlaw & De Vlieghere-He 1992; Kang et al., 2007). In our previously published model (Sarkar & Kundu 2025), the starting aperture size of the simulation during all three daytime phases was set to 7 µm. A basal value of 0.6 µm during all the six phases was considered. But we did not impose any upper limit on stomatal opening. In the present study, however, we allow the aperture size to increase up-to 7µm, above the basal stomatal opening of 0.6 µm, resulting in a maximum aperture size of 7.6 µm at any phase, which lies within the reported experimental range, mentioned earlier. As there was a majority of C_3_ plants on the Earth before Oligocene or Miocene period (Sage *et al*. 2023), we start the simulation using the constraints of C_3_ metabolism under the atmospheric condition of early-Eocene period with the average T of 35°C, RH of 70% and [CO_2_]_a_ of 800 ppm (**Figure 1b**). Next, we gradually decrease the [CO_2_]_a_, T and RH until the values reach to 250 ppm, 22°C and 30% respectively, representing the atmospheric condition of the early-Miocene period and simulate the model at each step (**Figures 1b, c**). The 2-step iteration method used for simulating the model at a single geological time point is given in **Figure 2a**.

**Figure 2.**
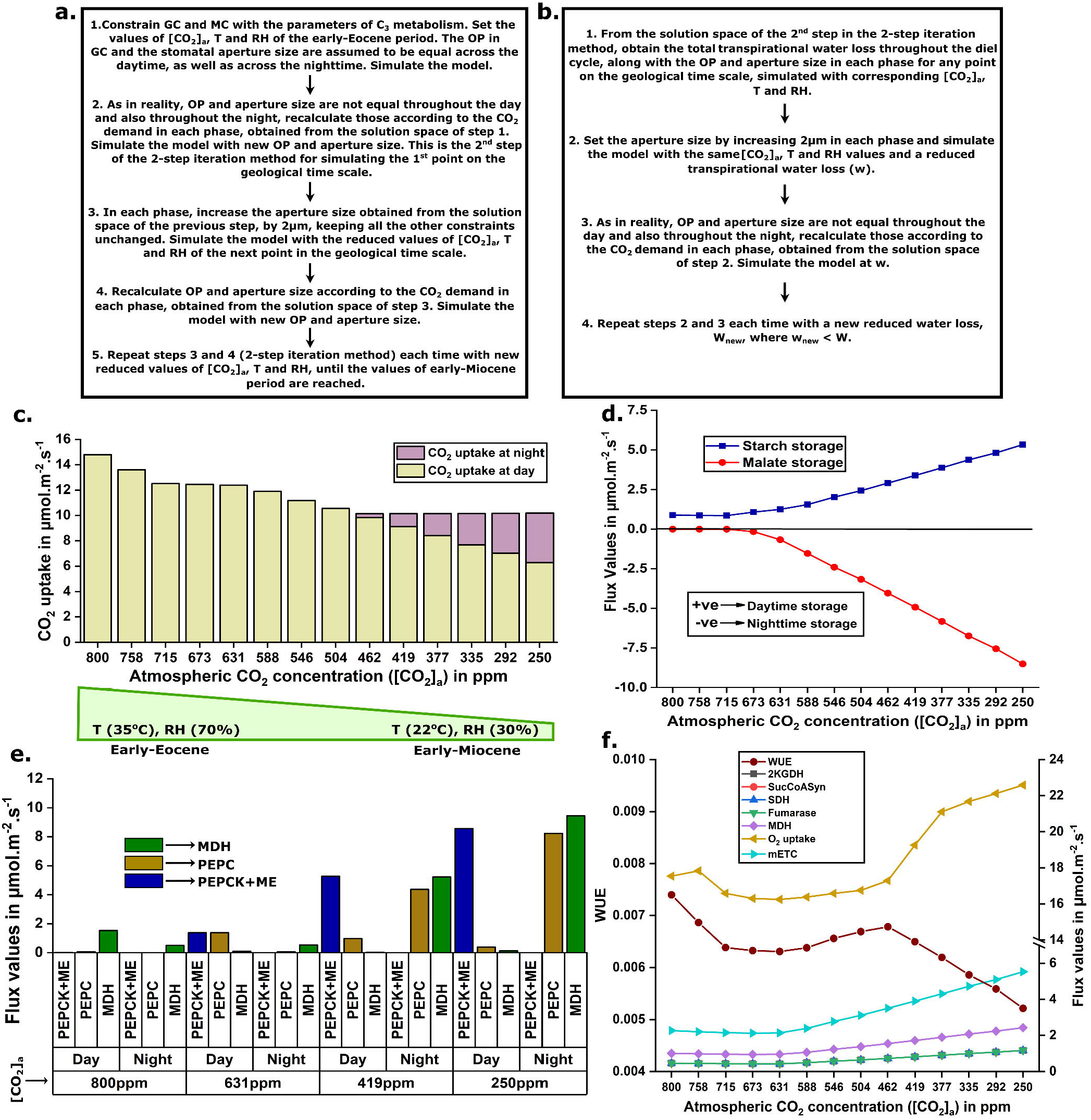
**a**. Steps for simulating the model under the changing atmospheric conditions between early-Eocene to early-Miocene time period. **b**. Steps for simulating the model with reducing transpirational water loss at any point of the atmospheric changes between that geological time interval. **c**. Variation in the total CO_2_ uptake throughout the diel cycle and the amount of CO_2_ uptake at day and night with the decrease in [CO_2_]_a_, T and RH. Total CO_2_ uptake declined due to refixation of respiratory CO_2_ (discussed in detail in main text). **d**. Changes in daytime starch storage and the nighttime malate storage with the decrease in [CO_2_]_a_, T and RH (declines in T and RH are shown in **c**). Increased storages of starch and malate at day and night respectively shows the C_3_-to-CAM transition. **e**. Changes in the activities of four C_4_-like enzymes related to carboxylation (PEPC and MDH) and decarboxylation (PEPCK and ME) of malate at night and day respectively with the decline in [CO_2_]_a_. Increased activities of PEPC, MDH at night and PEPCK, ME at day represent the metabolic shift towards CAM. **f**. Variation in the flux through mETC and 5 enzymes of TCA cycle in MC at night, total nocturnal O_2_ uptake and changes in WUE with the decrease in [CO_2_]_a_, T and RH (declines in T and RH are shown in **c**). Flux values of 2KGDH, SucCoASyn, SDH and fumarase overlap. Combinations of T and RH values corresponding to all the [CO_2_]_a_ values are listed in **Figure 1c**. It should be noted that all the flux values presented in the graphs are normalized by phloem sap production. 2KGDH, 2-ketogluterate dehydrogenase; SucCoASyn, succinyl-CoA synthetase; SDH, succinate dehydrogenase; mETC, mitochondrial electron transport chain; WUE, water use efficiency.

Moreover, to pinpoint the exact [CO_2_]_a_ at which the model initiates nocturnal CO_2_ uptake, we simulate the model across a fine gradient of [CO_2_]_a_, varying it with 1ppm increments between the points where nocturnal CO_2_ uptake is first observed and the immediately preceding point identified in the previous simulation (**Figure 2c**). We have also tested how the result outcomes depend on the variations in i) upper (in early-Eocene) and lower (in early-Miocene) values of RH and T and ii) difference of maximum and minimum values of T and RH throughout the diel cycle.

The increased aridity is expected to be associated with reduced water availability and this is also supported by the reports of disappearance of lakes caused by aridity (Wang et al., 2023) during various geological periods between early-Eocene and early-Miocene. As reduction in transpirational water loss is one of the ways to adapt and survive at water-stressed conditions, to check the effect of water scarcity at different geological time points, we simulate the model by gradually reducing transpirational water loss at every point of the atmospheric changes between early-Eocene to early-Miocene time period using the 2-step iteration method, shown in **Figure 2b**.

## 3. Results

### 3.1. C_3_-to-CAM transition under ancient atmospheric changes

At first, we simulate the model under the atmospheric condition of early-Eocene with [CO_2_]_a_ at 800 ppm, T at 35ºC and RH at 70%. Results confirm the characteristics of C_3_ metabolism (**Supporting Information Dataset S4**)-i) CO_2_ uptake only at daytime (Winter & Smith 2022), ii) malate storage at daytime in MC (Winter & Smith 2022), iii) early morning K^+^ storage in GC (Daloso *et al*. 2017) and sucrose storage in the later phases of the day in GC (Daloso *et al*. 2017) etc. Then, we gradually decrease the [CO_2_]_a_, T and RH until the values reach to 250 ppm, 22°C and 30% respectively, representing the atmospheric condition of the early-Miocene time period and simulate the model at each step, keeping the RuBisCO’s carboxylase-to-oxygenase activity ratio (C/O) fixed at 3 (experimentally observed value of C/O for C_3_ (Gutteridge & Pierce 2006)). With this gradual change in atmospheric conditions, the metabolism begins to exhibit key features of CAM (Winter & Smith 2022) (**Supporting Information Dataset S4**)-i) CO_2_ uptake at night through nighttime GC-mediated stomatal opening, ii) the increased activities of some C_4_-like enzymes such as PEPC and MDH at nighttime, whereas PEPCK and NADP-ME at daytime in MC; and iii) the increased daytime starch and nighttime malate storages in MC. Moreover, simulating the model with C/O fixed at 5.15:1 (experimentally observed values of C/O for CAM (Lüttge 2010)) under the ancient atmospheric condition of early-Eocene showed that the ratio of 5.15:1 alone does not induce the CAM pathway (**Supporting Information Dataset S4**). These simulations indicate that increasing the C/O alone does not drive CAM evolution; instead, the C_3_-to-CAM transition is primarily driven by the ancient atmospheric changes. Therefore, to incorporate the known physiological conditions, we simulate the model under varying ancient atmospheric conditions with gradually increasing the C/O from 3:1 to 5.15:1 (also used in previous computational studies of C_3_-to-CAM transition (Tay *et al*. 2021; Sarkar & Kundu 2025)) and observe the emergence of previously mentioned traits of CAM (**Figures 2c-e**, (**Supporting Information Dataset S4**)). In CAM, this daytime stored starch is broken down via glycolysis at night to generate phosphoenolpyruvate (PEP), the substrate for the carboxylating enzyme, PEPC. PEPC then fixes both the atmospheric and nocturnal respiratory CO_2_ (Griffiths et al. 1989; Lim et al., 2019; Gilman et al. 2023; Winter & Holtum 2024) in a four-carbon compound, OAA which is subsequently converted to malate by cytosolic MDH. This malate is stored overnight in vacuole and decarboxylated at daytime by the decarboxylating enzymes PEPCK and ME, releasing CO_2_ for fixation in CBB cycle. Although nocturnal CO_2_ uptake begins around 483 ppm, very close to a previous prediction of CAM diversification (Hönisch *et al*. 2023), nighttime malate accumulation starts earlier at about 673 ppm (**Figure 2d**). In response to decline in [CO_2_]_a_, plant initially adjusts its metabolism by limiting daytime CO_2_ uptake and refixing respiratory CO_2_ into malate at night (**Figure 2c**) and later, starts taking atmospheric CO_2_ at night to meet the total carbon requirement, in addition to respiratory CO_2_ fixation (Griffiths *et al*. 1989). Results further show that nocturnal mitochondrial respiration increases with the emergence of CAM. This is supported by elevated nocturnal O_2_ uptake, and increased flux values through several TCA cycle enzymes including 2-ketogluterate dehydrogenase (2KGDH), succinyl-CoA synthetase (SucCoASyn), succinate dehydrogenase (SDH), fumarase, and malate dehydrogenase (MDH) (**Figure 2f, Supporting Information Dataset S4**). Among these, 2KGDH, SDH and MDH produce NADH and FADH_2_, which subsequently participate in mitochondrial electron transport chain (mETC) to produce higher amount of ATP. This increased production of mitochondrial ATP, essential for CAM cycle, also enhances the demand for the nocturnal O_2_ uptake. Additionally, simulation with restricted nighttime O_2_ uptake reveals that increased nocturnal O_2_ uptake is mandatory for CAM functioning, supporting the experimental observation that CAM requires elevated nocturnal respiratory rate and nocturnal O_2_ consumption (Leverett & Borland 2023). Furthermore, as we move towards Miocene, there is an overall decrease in water-use efficiency (WUE) (**Figure 2f**) due to increase in transpirational water loss. This result is consistent with earlier experimental reports suggesting that plants cope with the lower [CO_2_]_a_ by opening stomata and therefore stomatal conductance increases (Cowan & Farquhar 1977; Sage *et al*. 2023). This consequently increases the transpiration rate and hence WUE decreases (Sage *et al*. 2023).

Now, to identify the major atmospheric changes that favour C_3_-to-CAM transition, we study the response of plant C_3_ metabolism at higher [CO_2_]_a_ (800 ppm), but at reducing T (35ºC to 22 ºC) and RH (70% to 30%) (**Supporting Information Dataset S5**). Interestingly, we observe variations of T and RH, alone or in combination, are not sufficient to drive the C_3_-to-CAM transition. We already showed that the transition is favoured when changes in T and RH are coupled with lower [CO_2_]_a_ (**Figures 2c-e**). Moreover, when we simulate the C_3_ metabolism at reducing [CO_2_]_a_ only, keeping the T and RH fixed at 35ºC and 70% respectively, we observe the C_3_-to-CAM transition (**Supporting Information Dataset S6**). These support the hypothesis that lower [CO_2_]_a_ during ancient times drives the C_3_-to-CAM transition (Sage *et al*. 2023).

### 3.2. CAM emergence with decreasing water availability or increasing aridity even with high [CO_2_]_a_

In the previous simulations, we have not restricted the transpirational water loss to nullify the effect of reduced water loss in C_3_-to-CAM transition established by previous studies (Töpfer *et al*. 2020; Sarkar & Kundu 2025). However, the increased aridity of early-Miocene is expected to be associated with reduced water availability and this water scarcity is also supported by the disappearance of lakes caused by aridity (Dupont-Nivet *et al*. 2007; Liu *et al*. 2014; Wang *et al*. 2023) during various geological periods between the Eocene and Miocene. As reduction in transpirational water loss is one of the ways to adapt and survive at water-stressed conditions, we gradually reduce the transpirational water loss at different points between early-Eocene to early-Miocene (**Figure 2b**) and observe a metabolic shift to CAM regardless of whether [CO_2_]_a_ is high or low (**Supporting Information Datasets S4**). We also observe the shift in stomatal opening from day to night (**Figure 3e, Supporting Information Figure S2**), maximum CO_2_ uptake at night (**Figure 3a,b Supporting Information Figure S3**), higher activities of PEPC, PEPCK, MDH and NADP-ME (**Figure 3c,d, Supporting Information Figure S4**), and increased storages of starch and malate at day and night respectively (**Supporting Information Figure S5**). These findings support the fact that water stress shifts the C_3_ metabolism to CAM even at higher CO_2_ (Males & Griffiths 2017). Moreover, for any given [CO_2_]_a_, a reduction in the transpirational water loss forces the model to decrease the phloem sap production, clearly revealing a trade-off between water conservation and phloem sap production. It should be noted that all the flux values presented in the graphs (**Figures 2** and **3**) are normalized by phloem sap production. Interestingly, WUE increases when associated with the limitation in transpirational water loss at any point between early-Eocene and early-Miocene (**Supporting Information Figure S6**). As reduction in transpirational water loss (or water-saving) diminishes with decline in [CO_2_]_a_, the extent of WUE improvement due to reduced water loss, is also reduced.

**Figure 3.**
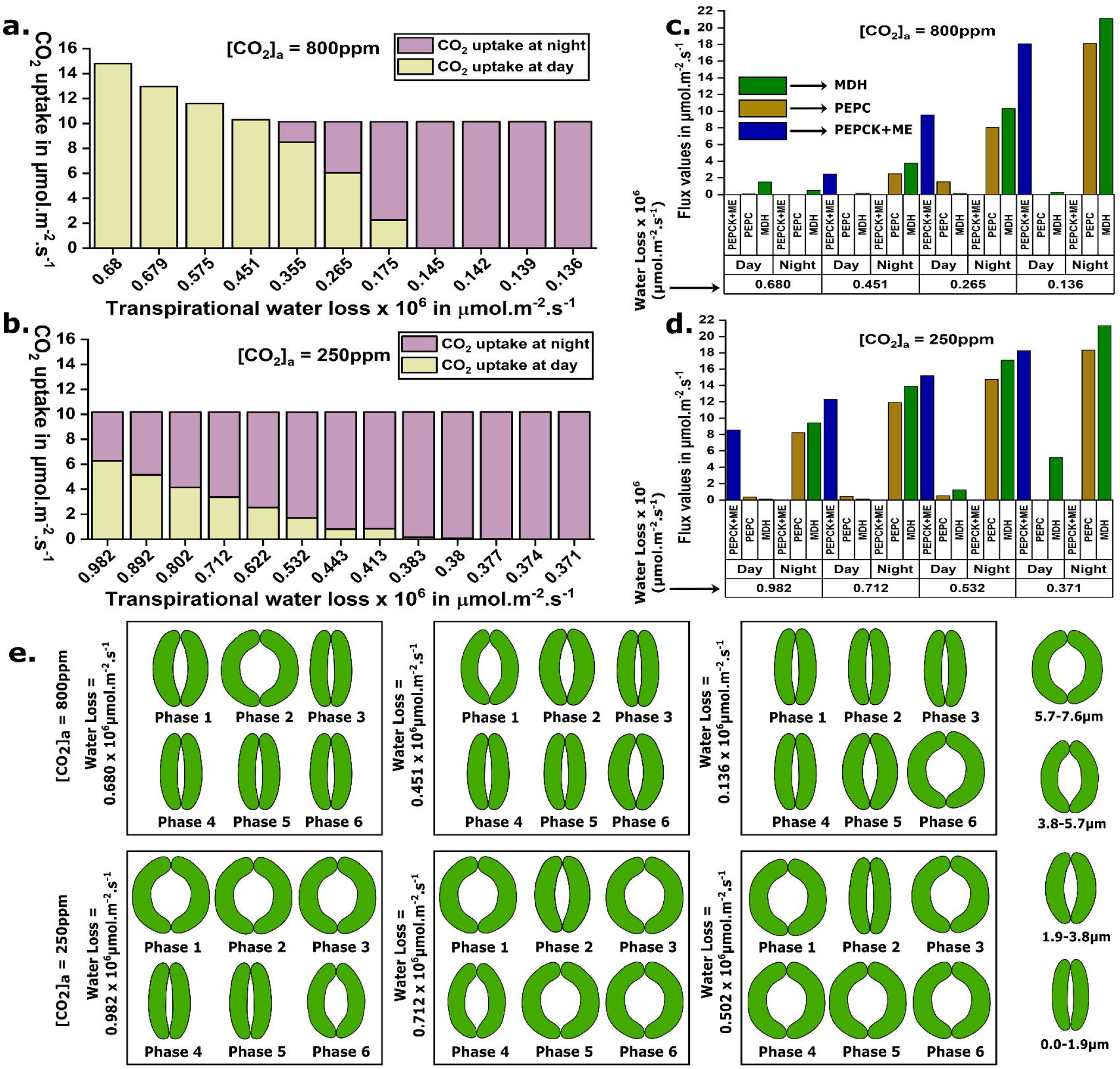
**a-b** show the variation in CO_2_ uptake at day and night with reduction in transpirational water loss at different [CO_2_]_a_. **c-d** show the changes in the activities of the carboxylating (PEPC and MDH) and decarboxylating (PEPCK and ME) enzymes with reduced transpirational water loss at different [CO_2_]_a_. **e**. Variations in stomatal aperture sizes at day and night with decline in [CO_2_]_a_ (row-wise) and transpirational water loss (column-wise) are shown. Reduced [CO_2_]_a_ forces the nighttime stomatal opening along with day, however, at a fixed [CO_2_]_a_, stomatal opening shifts from day to night with the decrease in transpirational water loss. Only [CO_2_]_a_ values are shown in the graphs (a-e). Combinations of T and RH values corresponding to all the [CO_2_]_a_ values are listed in **Figure 1c**. It should be noted that all the flux values presented in the graphs are normalized by phloem sap production.

### 3.3. Changes in enzymatic activities in MC and GC during C_3_-to-CAM transition

While the fluxes through many reactions of central carbon metabolism show temporal changes during this transition, we expect there might be a gradual increase/decrease for some of them. To identify these enzymatic shifts during the transition, we calculate the Spearman’s rank correlation between the flux values through each of the enzymatic reactions and intracellular transporters across different cell types and the six phases of the diel cycle with-i) the gradual atmospheric changes during early-Eocene to the early-Miocene and ii) the gradually reduced transpirational water loss at each point between the same geological time interval (**Supporting Information Dataset S7**). The correlation values reveal a similar trend in the shift of the temporally differential flux distribution pattern in MC during the transition in both the cases. This pattern also aligns with our earlier findings during the C_3_-to-CAM transition in reduced transpirational water loss at a fixed T, RH and [CO□]□ (Sarkar & Kundu 2025). The enzymes showing gradual changes in MC are spanned across various metabolic pathways, including carboxylation-decarboxylation, glycolysis, gluconeogenesis, TCA cycle, pentose phosphate pathway, mitochondrial ETC etc. However, GC exhibits different patterns, possibly due to the different stomatal behaviours. Under condition (i), as [CO_2_]_a_ declines, the plant’s CO_2_ demand necessitates wider stomatal opening (**Figure 3e, Supporting Information Figure S2**). As the stomatal pore has an upper physiological limit, very low [CO_2_]_a_ forces the plant to open their stomata both at day and night to meet the required carbon demand (**Figure 3e, Supporting Information Figure S2**). However, under condition (ii), at higher [CO_2_]_a_ (800 ppm), the minimization of transpirational water loss mainly drives the C_3_-to-CAM transition, and a large number of reactions in GC show similar pattern as observed in our previous study (Sarkar & Kundu 2025). In contrast, at lower [CO_2_]_a_ (250 ppm), CO_2_ limitation alone significantly drives CAM-like features even without any restriction in water loss, and further reduction in water loss has minimal impact, resulting in lower water-saving.

### 3.4. Simulations at extended model also show both [CO_2_]_a_ decline and aridity favour C_3_-to-CAM transition

Research suggests that species differ in how stomatal density responds to [CO_2_]_a_-some increasing, some decreasing, and others showing none or only minor changes (Serna, 2008). We now aim to examine how the outcomes of our simulations depend on these different stomatal density responses under varying [CO_2_]_a_. As mentioned, several species exhibit no change in stomatal density in response to elevated [CO_2_]_a_ (Oberbauer et al. 1985; Radoglou & Jarvis 1990, 1992; Ferris & Taylor 1994; Woodward & Kelly 1995; Apple et al. 2000; Serna & Fenoll 2000). All the previous results presented in this study are based on simulations assuming constant stomatal density across changing [CO_2_]_a_. In some other plants, stomatal density increases with declining [CO_2_]_a_ (Woodward 1987; Woodward & Bazzaz 1988; Woodward & Kelly 1995). To incorporate this into our analysis, we simulate the model by gradually increasing stomatal density from 250 mm^-2^ as [CO_2_]_a_ declined. Results show that higher stomatal density allows greater daytime CO_2_ uptake at each [CO_2_]_a_ level compared with our previous results, leading to a delayed transition from C_3_ to CAM metabolism in the ancient time scale (**Supporting Information Dataset S8**). In some other species, stomatal density decreases with the decline in [CO_2_]_a_(Ferris & Taylor 1994; Woodward & Kelly 1995; Driscoll et al. 2006). To evaluate this scenario, we simulated our model by gradually decreasing stomatal density from 250 mm^-2^ as [CO_2_]_a_ declined. The results show that lower stomatal density reduces daytime CO_2_ availability compared with our previous simulations, causing the shift from C_3_ to CAM metabolism to occur earlier in the geological time scale (**Supporting Information Dataset S8**).

It should be noted that RH values for the early-Eocene and early-Miocene are not available, whereas the middle Eocene RH has been estimated to be 67% (Jahren *et al*. 2003). Several studies indicate that RH declined over this time interval (Passchier *et al*. 2013; Wang *et al*. 2016). Therefore, initially we have used RH values of 70% for the early-Eocene and 30% for the early-Miocene, representing conditions slightly higher and lower than those of the middle Eocene, respectively. However, due to the lack of precise RH estimates for these ancient periods, we have also conducted simulations using different RH values of early-Eocene and early-Miocene to test how changes in RH influence the model outcomes. The results (Data is not shown, however, the model and code are provided in **Supporting Information Datasets S1-S3**) indicate that even with a variation in the upper (RH during early-Eocene) and lower (RH during early-Miocene) bounds of RH, the model consistently predicts a transition from C_3_ to CAM metabolism. Although transpirational water loss changes in response to RH fluctuations, the overall conclusions of the study remain unchanged.

In our model, K_2_malate, KCl and sucrose are initially considered as osmolytes (Talbott & Zeiger 1996; Lawson 2009; Daloso et al., 2016; Daloso et al. 2017; Santelia & Lawson 2016; Robaina-Estévez et al.,2017), with only sucrose permitted to transfer from MC to GC (Lawson, Simkin, Kelly & Granot 2014; Daloso *et al*. 2016). We now extend the model to include additional osmolytes (nitrate, glucose, fructose and maltose) in GC and transfer metabolites (glucose and malate) from MC to GC (Talbott & Zeiger 1993; Lawson et al. 2014; Daloso et al. 2017; Flütsch et al. 2020b a; Dang et al., 2024) (**Supporting Information Dataset S9**). Simulations of this extended model also indicate that, even with these inclusions, declining [CO_2_]_a_ and rising aridity consistently drive a metabolic shift toward CAM. While, MC shows similar metabolic patterns, GC shows variation in the storage patterns of these osmolytes for generating turgor pressure (**Supporting Information Dataset S10**).

### 3.5. Predicting the effect of atmospheric change in future

To understand the effect of changing atmospheric conditions of future, we further simulate the model under varying [CO_2_]_a_, T, and RH and observe-i) metabolism shifts towards C_3_ under elevated [CO_2_]_a_ (**Supporting Information Table S2**), ii) water loss increases with higher T, lower [CO_2_]_a_, and reduced RH (**Supporting Information Figure S7a, Supporting Information Dataset S11**) and iii) CAM is induced with greater nocturnal CO_2_ uptake (**Supporting Information Figure S7b, Supporting Information Dataset S11**) for a fixed T, [CO_2_]_a_ and water loss with reduced RH (for details, see **Supporting Information File**).

## 4. Discussion

Simulating life– or replicating cellular and metabolic processes to trace potential evolutionary changes under ancient Earth conditions– is one of the most challenging and valuable areas of scientific research (Miller 1953). At the same time, we are facing severe existential threats from global climate change and extreme weather events, which pose additional challenges such as ensuring food security (Lim *et al*. 2025). Since plants with CAM-like metabolism perform well in arid environments in terms of survival and water use, one of the possible solutions is to bioengineer CAM-like metabolism into C_3_ plants to develop drought-resistant crop varieties with improved water use efficiency. It necessitates significant physiological, anatomical, as well as metabolic modifications within C_3_ plants. Several phylogenomic and large-scale carbon isotope studies suggest that CAM evolved from the C_3_ photosynthesis approximately 20-30 MYA and atmospheric changes during that period favoured the metabolic transition from C_3_ to CAM (Heyduk 2022; Sage *et al*. 2023). To test this hypothesis, one would need to recreate ancient atmospheric conditions, place a C_3_ plant in that environment, and observe it over time to detect any metabolic transition. Challenges lie in setting up the necessary experimental arrangements as well as a reasonable time of observation. Although several experimental studies examined different aspects of C_3_-to-CAM transitions (Tan & Chen 2023) (for details, see the references therein), most of the data are for entire daytime or nighttime period and the experiments were not performed under ancient conditions. On the other hand, while most of the previous computational efforts are limited to investigate the transition in MC only (Töpfer *et al*. 2020; Tay *et al*. 2021), our recent study on an environment coupled metabolic model captures potential metabolic transitions under water-saving conditions (Sarkar & Kundu 2025), but does not account for atmospheric changes during ancient times.

Here, we have used a refined version of our previously developed bio-physicochemical model that integrates cellular metabolism, gaseous exchange between the environment and the leaf’s internal air space, and transpirational water loss which depends on the amount of CO_2_ uptake, the diel fluctuations in T and RH, and the difference between internal and external CO_2_ concentrations. We have reconstructed ancient environmental conditions over millions of years by gradually varying T, RH and [CO_2_]_a_ from early-Eocene to early-Miocene and simulated the model under this changing atmospheric conditions. It is important to note that this study does not model the carbon-concentrating mechanism of CAM in terms of concentrating CO_2_ around RuBisCO and restricting it to interact with O_2_; rather, it focuses on differential metabolic changes in terms of flux distributions and inverted stomatal behaviour during the C_3_-to-CAM transition. Results show that changing C/O from 3 to 5.15 alone cannot favour the transition, rather changing atmosphere (decreasing T, RH and [CO_2_]_a_ from early-Eocene to early-Miocene) can do. Therefore, while studying the effect of ancient atmospheric changes to get the differential flux distributions through metabolic pathways, CO2 uptake, osmolyte accumulation dependent stomatal rhythm, we gradually increase C/O from 3 (experimentally observed value for C_3_ (Gutteridge & Pierce 2006)) to 5.15 (experimentally observed value for CAM (Lüttge 2010)) to use a more realistic constraint on RuBisCO’s C/O activity, also used in previous studies (Sarkar & Kundu 2025; Tay et al. 2021). We also observe that changes in T or RH alone, or in different combinations, could not induce CAM-like metabolism, whereas a reduction in [CO_2_]_a_ alone with a fixed T and RH is capable of initiating the metabolic shift. This suggests that declining [CO_2_]_a_ could have favoured the evolution of CAM during the ancient times. It is also important to note that as [CO_2_]_a_ declines, the amount of CO_2_ entering the leaf decreases for a fixed stomatal aperture. When [CO_2_]_a_ becomes sufficiently low such that the CO_2_ entering the leaf even with fully open stomata throughout the day is insufficient, we observe beginning of nocturnal CO_2_ uptake probably to produce required amount of phloem sap, needed for plant. Thus, although the metabolism shifts towards CAM with declining [CO_2_]_a_, the completely inverted stomatal pattern, an important characteristic of CAM, is not observed (Sarkar & Kundu 2025). The inverted stomatal pattern is observed when we restrict the transpirational water loss. In fact, geological data also suggest that the disappearance of lakes at various time points during this period (Dupont-Nivet *et al*. 2007; Wang *et al*. 2023). While declining [CO_2_]_a_ leads to increased nighttime PEPC activity, enhanced daytime starch and nighttime malate accumulation, and nocturnal CO2 uptake, it alone does not induce daytime stomatal closure, as mentioned earlier. In contrast, reducing transpirational water loss reproduces all the features of CAM observed under declining [CO_2_]_a_ and additionally promotes daytime stomatal closure. We further observe gradual changes in flux through the reactions involved in different pathways of central carbon metabolism such as glycolysis, gluconeogenesis, TCA cycle, ETC, PPP, different shuttles between different compartments etc., in MC in both the cases, but the patterns differ in GC (**Supporting Information File**). This suggests that declining [CO_2_]_a_ and increasing aridity may exert slightly different effects on osmolytes accumulation dependent stomatal opening and also indicates the flexibility in selecting osmolytes to accumulate for stomatal opening. In recent years, multiple theories have been proposed and debated regarding osmolyte accumulation in GC across different phases (Granot & Kelly 2019). Earlier studies suggested that stomatal opening in C_3_ plants is primarily driven by the accumulation of K+ and its counterions during the early morning, while sucrose plays a major role in maintaining stomatal opening during the later phases of the day (Talbott & Zeiger 1996; Lawson 2009; Santelia & Lawson 2016; Daloso *et al*. 2017). However, a recent study in *Vicia faba* indicates that glucose also contributes significantly to early-morning stomatal opening in C_3_ plants (Flütsch et al. 2020b, a). Present study shows that in C_3_, K+, glucose, and fructose contribute to stomatal opening in phase 1, whereas in phase 2, glucose and fructose majorly balances the OP. In the last phase of the day, maltose accumulates and restricting it leads to increased accumulation of sucrose to maintain the OP. In addition, accumulation of NO3- and citrate3-is also observed when the accumulation of other metabolites is restricted. Our study also reveals phase-dependent accumulation of different osmolytes across different phases of CAM, which has also been experimentally reported in CAM plants, like *Kalanchoë fedtschenkoi* (Abraham *et al*. 2020; Hurtado-Castano *et al*. 2023). Moreover, enhanced activities of enzymes involved in TCA cycle and mETC, along with increased ATP production reflect increased respiration in CAM. Our study further shows that restricting nocturnal O_2_ uptake prevents the metabolism from shifting to CAM, indicating the importance of nocturnal O_2_ uptake and enhanced respiration in the evolution of CAM. Interestingly, atmospheric O_2_ concentration increased during that geological time period (Berner et al. 2007), which probably supports the higher O_2_ consumption and facilitated the emergence of CAM.

Here, we have used the maximization of phloem sap production as one of the objectives of simulation. The results show that when the metabolism shifts toward CAM under changing atmospheric conditions, phloem sap production is reduced by only ∼5%. In contrast, when CAM emerges due to restrictions on transpirational water loss, phloem sap production decreases by approximately 50% across any point between the early Eocene and early Miocene. These findings suggest a trade-off between water-saving and phloem sap production, which has also been reported in previous studies (Töpfer *et al*. 2020).

Furthermore, it has been experimentally reported that stomatal density responds differently to [CO_2_]_a_ across plant species. Stomatal density can increase or decrease or remain constant with decreasing [CO_2_]_a_ (Serna 2008). When stomatal density was gradually increased or decreased in our simulations during the transition in ancient atmospheric conditions, similar results were obtained; only the timing of nighttime CO2 uptake shifted along the geological time scale. Likewise, varying the upper (early-Eocene) and lower (early-Miocene) values of T and RH did not significantly affect the conclusions drawn from the simulations.

Although we observe a metabolic shift toward CAM under the changing atmospheric conditions of ancient times, transpirational water loss increases when no restriction is imposed. During this geological period, RH declined, and evidence such as the disappearance of lakes with increasing aridity during ancient times suggests reduced water availability (Dupont-Nivet *et al*. 2007; Wang *et al*. 2023). Therefore, to account for this constraint, we restricted transpirational water loss during the transition toward the early-Miocene, keeping it at the value observed under the higher RH conditions of the early-Eocene. To examine this effect, we performed an additional simulation. First, we calculated the transpirational water loss in the early Eocene (denoted as *w*). Keeping this water loss fixed at *w*, we then simulated the transition toward the early Miocene by gradually reducing atmospheric CO_2_ concentration ([CO□]_a_), T, and RH, as described in subsection 3.1. The results show that additional restriction on transpirational water loss, as a consequence of increasing aridity, promotes the emergence of stronger CAM characteristics. Nighttime CO2 uptake begins earlier along the geological timeline, and by the early Miocene, the majority of CO2 uptake occurs at night. Additionally, the combined effects of declining [CO_2_]_a_ and restricted water loss lead to the emergence of an inverted stomatal rhythm. These findings suggest that the combined effects of both the evolutionary drivers-declining [CO_2_]_a_ and increasing aridity could have driven the evolution of CAM. Future [CO_2_]_a_ increase might favour C_3_ over CAM; but aridity or water-scarcity will favour CAM.

## 5. Conclusion

To summarize, although anatomical and physiological traits such as increased succulence in CAM plants (Lüttge 2004), stomatal regulation by differential activities of signalling pathways (Blatt 2000), mechanistic regulation of stomata (Blatt et al. 2022), change in assimilation rate with atmospheric conditions (Rudenko *et al*. 2022) etc., are not included in our model; this environment-coupled simple cellular metabolic model-i) successfully reconstructs ancient conditions, ii) captures the C_3_-to-CAM transition driven by environmental changes during that geological time period, iii) successfully predicts known biochemical changes during C_3_-to-CAM transition, and iv) provides an initial useful framework to predict the possible differentially active metabolic pathways and quantitative estimations of relative flux values in the changing environment, a requisite for proper designing of abiotic stress tolerant cultivar.

## Acknowledgements

DS thanks Department of Biotechnology, Government of India for her fellowship.

