## Supporting Information File for "Ancient atmospheric changes could favour C_3_-to-CAM transition: A simulation-based evidence"

**Supporting Information Dataset S1.** The model file in excel format.

**Supporting Information Dataset S2.** The model file in sbml format.

**Supporting Information Dataset S3.** Python scripts for all the simulations.

**Supporting Information Dataset S4.** Flux solutions i) obtained from simulating the model by gradually reducing [CO_2_]_a_, T and RH, keeping the RuBisCO’s carboxylase-to-oxygenase activity ratio (C/O) fixed at 3, ii) obtained from simulating the model by gradually reducing [CO_2_]_a_, T and RH, keeping the C/O fixed at 5.15, iii) for each step of C_3_-to-CAM transition due to changes in atmospheric conditions between early-Eocene to early-Miocene with gradually increasing C/O from 3 to 5.15, iv) for each point of the transition due to reduced transpirational water loss studied at each step of the atmospheric changes and v) for each step of C3 to CAM transition due to changes in atmospheric conditions (reduced [CO2]a, T and RH) between early-Eocene to early-Miocene, keeping the water loss fixed at the value of transpirational water loss obtained in early-Eocene (1st point of the transition).

**Supporting Information Dataset S5.** Flux solutions obtained from simulating the model with reducing T (35-22°C) and RH (70-30%) at a fixed [CO_2_]_a_ of 800 ppm.

**Supporting Information Dataset S6.** Flux solutions obtained from simulating the model under different reduced [CO_2_]_a_ values at fixed T (35°C) and RH (70%).

**Supporting Information Dataset S7.** Correlation values of each reaction i) with change in atmospheric conditions (reduced [CO_2_]_a_, T and RH) between early-Eocene to early-Miocene and ii) with reduced transpirational water loss at each step of the atmospheric changes between that geological time interval.

**Supporting Information Dataset S8.** Flux solutions obtained from simulating the model with increasing and decreasing stomatal density in each step of C_3_-to-CAM transition due to changes in atmospheric conditions between early-Eocene to early-Miocene.

**Supporting Information Dataset S9.** The model file including additional osmolytes (nitrate, glucose, fructose and maltose) and transfer metabolites (glucose and malate) in excel format.

**Supporting Information Dataset S10.** Flux solutions obtained from simulating the model with additional osmolytes and transfer metabolites i) for each step of C_3_-to-CAM transition due to changes in atmospheric conditions between early-Eocene to early-Miocene and ii) for each point of the transition due to reduced transpirational water loss studied at each step of the atmospheric changes.

**Supporting Information Dataset S11.** Flux solutions obtained from simulating the model by reducing RH from 80% to 40% with different combinations of T (26-35°C) and [CO_2_]_a_ (800, 419 and 250 ppm), either keeping the water loss free or keeping the water loss fixed at a value obtained with 80% RH for the corresponding T and [CO_2_]_a_ values.

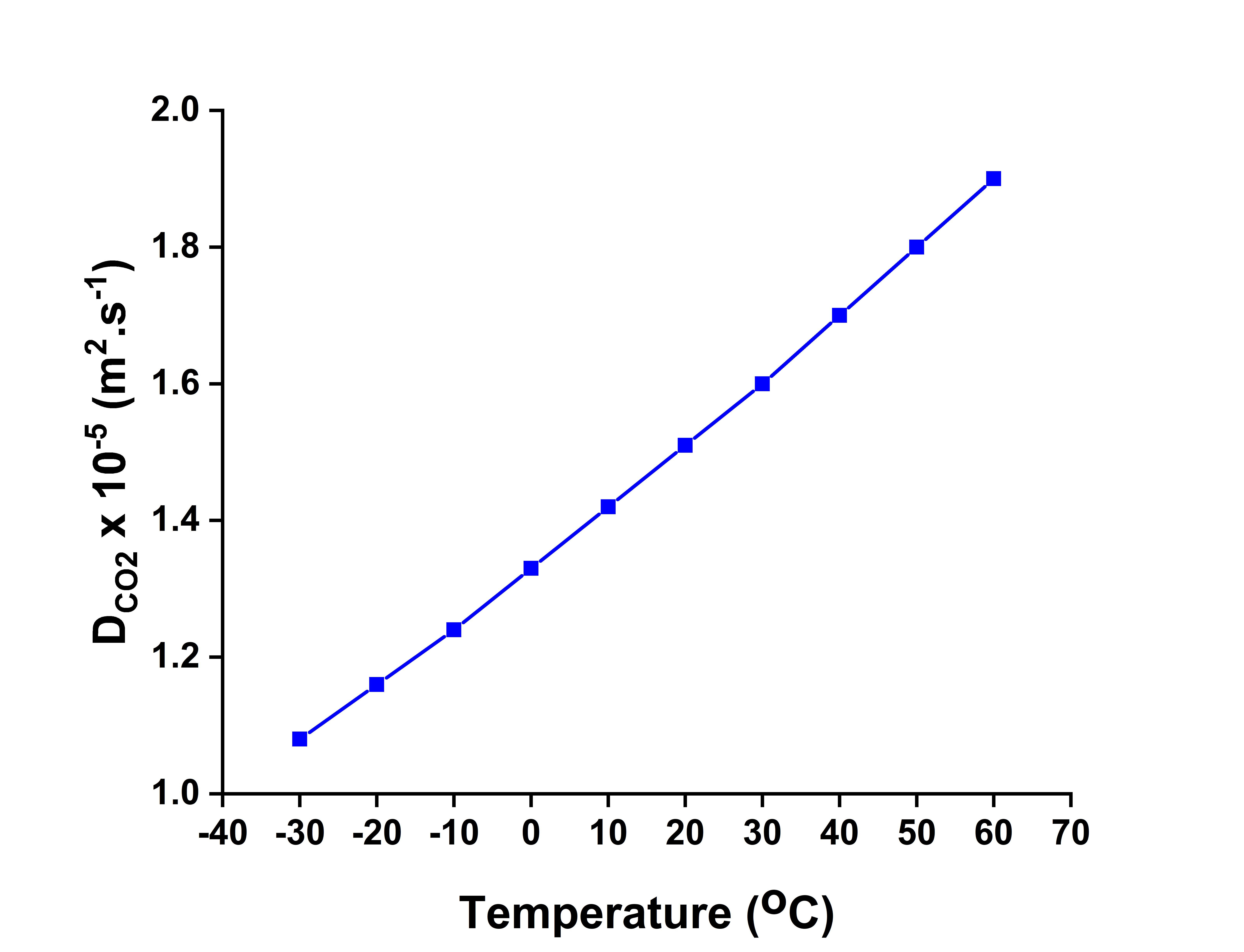

**Supporting Information Figure S1.** **Variation in the diffusion coefficient of CO_2_ (**$\mathbf{D}_{\mathbf{CO}_{\mathbf{2}}}$**) with temperature (T).**

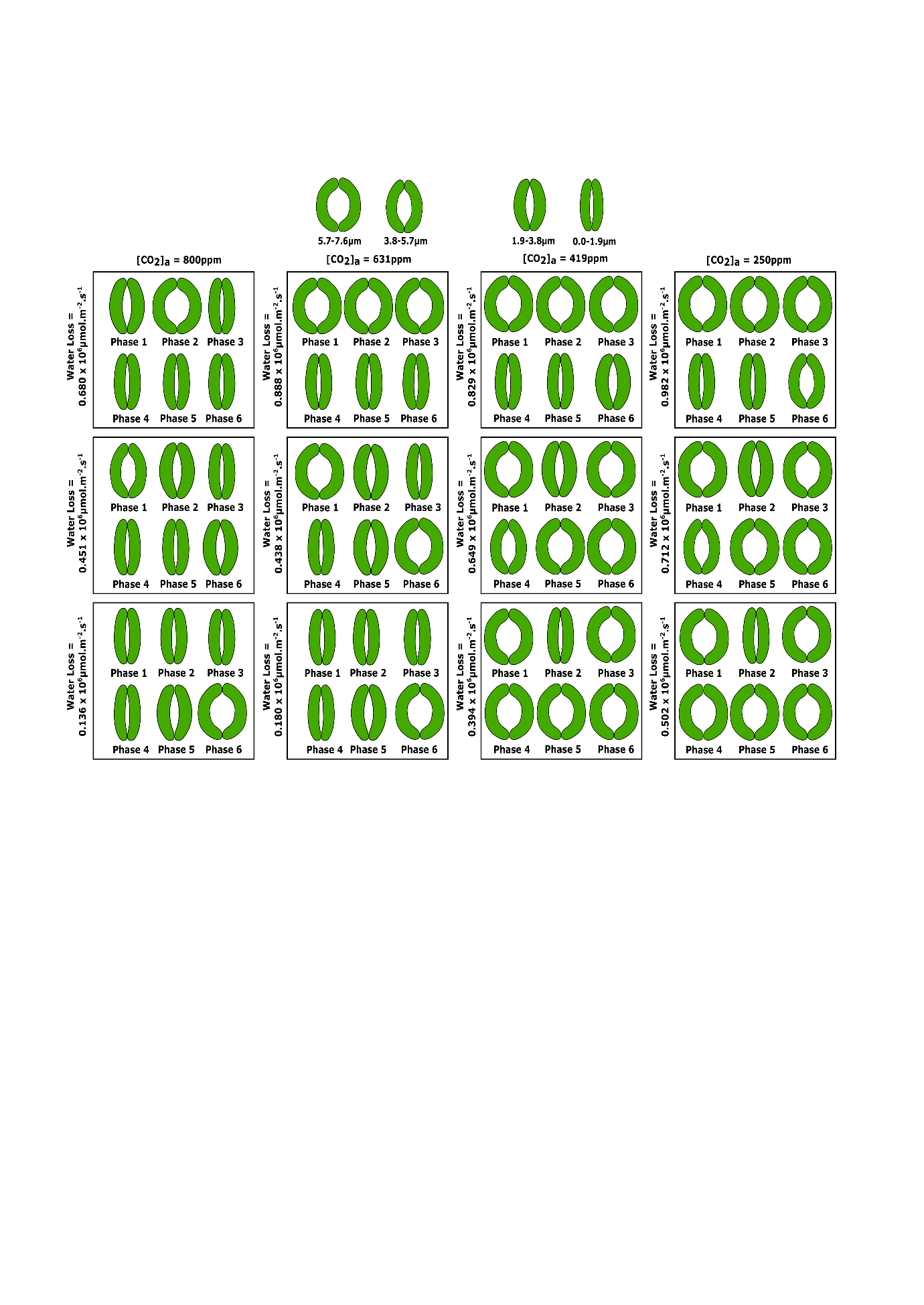

**Supporting Information Figure S2. Change in stomatal aperture size in different phases of day and night.** Variations in stomatal aperture sizes at day (phases 1-3) and night (phases 4-6) with i) decline in [CO_2_]_a_ (row-wise) and ii) decline in transpirational water loss (column-wise) are shown. Only [CO_2_]_a_ values are shown in all the graphs. Corresponding T and RH values are listed in **Supporting Information Fig. S1c**.

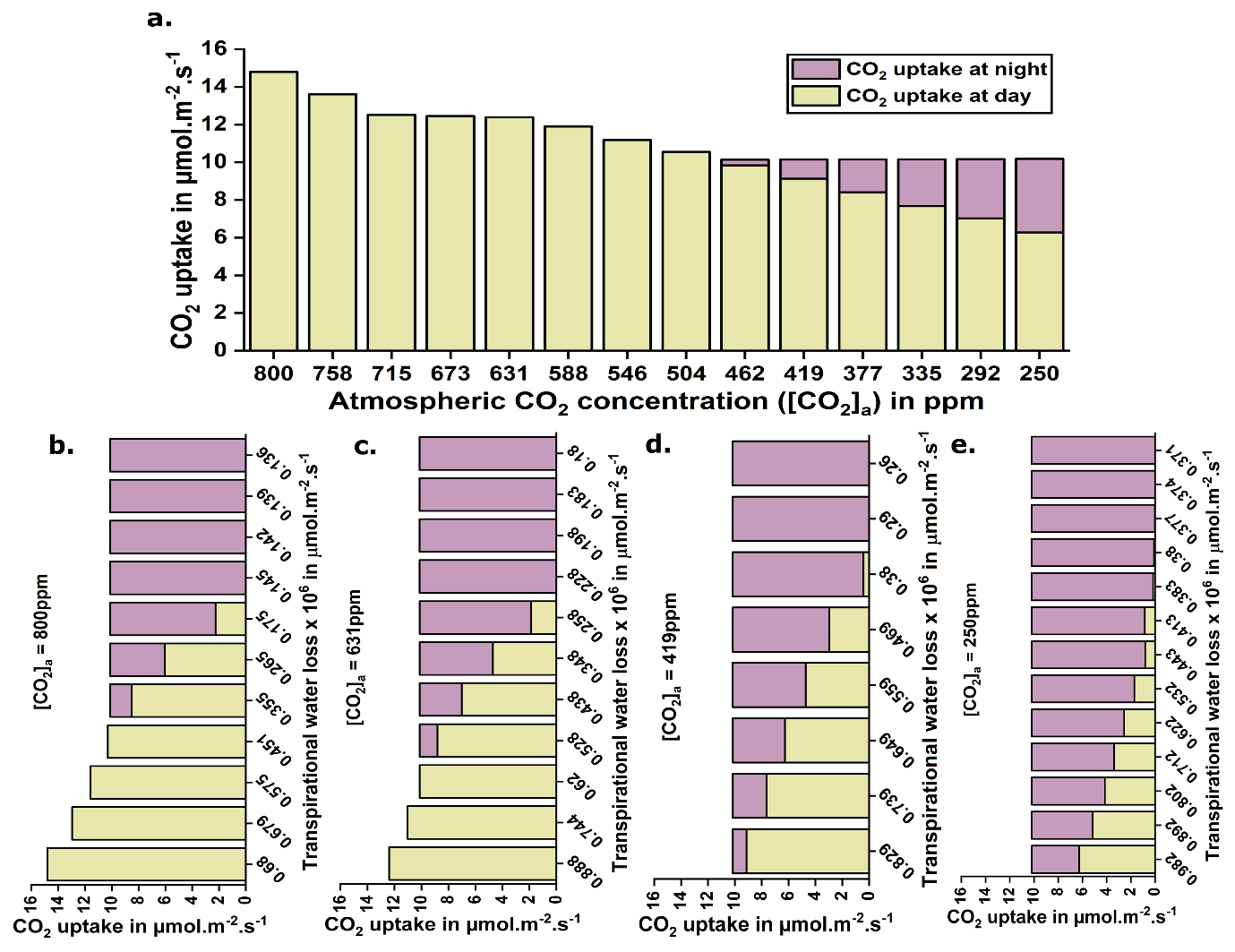

**Supporting Information Figure S3. Changes in the CO_2_ uptake at day and night. a** shows the variation in CO_2_ uptake at day and night with decreasing [CO_2_]_a_; and **b-e** show the variation in CO_2_ uptake at day and night with decreasing transpirational water loss at different [CO_2_]_a_. Only [CO_2_]_a_ values are shown in all the graphs. Corresponding T and RH values are listed in **Supporting Information Fig. S1c**. It should be noted that all the flux values presented in the graphs are normalized by phloem sap production.

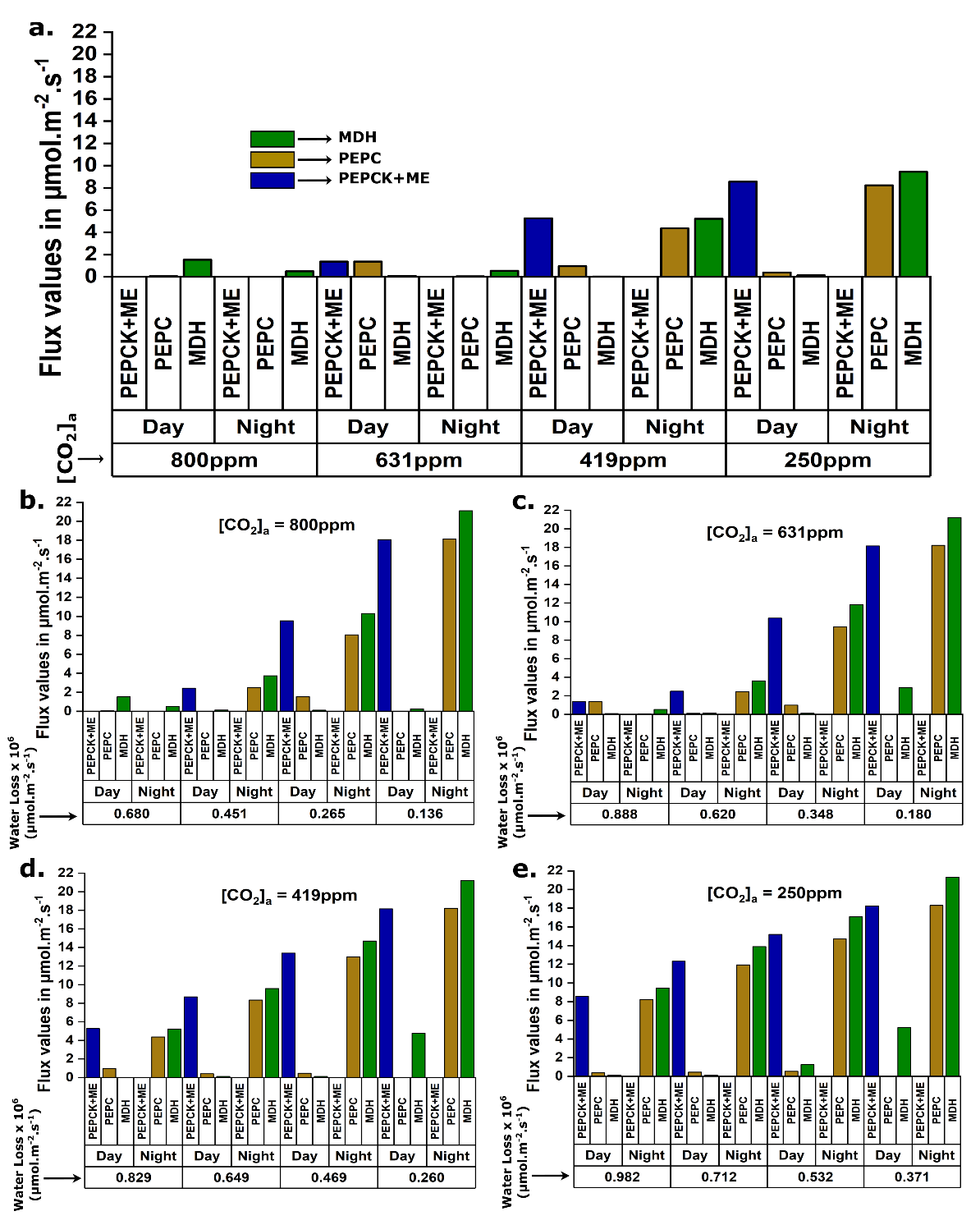

**Supporting Information Figure S4. Variations in flux values of different carboxylating and decarboxylating enzymes. a** shows changes in the activities of four C_4_-like enzymes related to carboxylation (PEPC and MDH) and decarboxylation (PEPCK and ME) of malate at night and day respectively with the decline in [CO_2_]_a_; and **b-e** show the changes of the activities of the same enzymes with decreasing transpirational water loss at different [CO_2_]_a_. Only [CO_2_]_a_ values are shown in all the graphs. Corresponding T and RH values are listed in **Supporting Information Fig. S1c**. It should be noted that all the flux values presented in the graphs are normalized by phloem sap production.

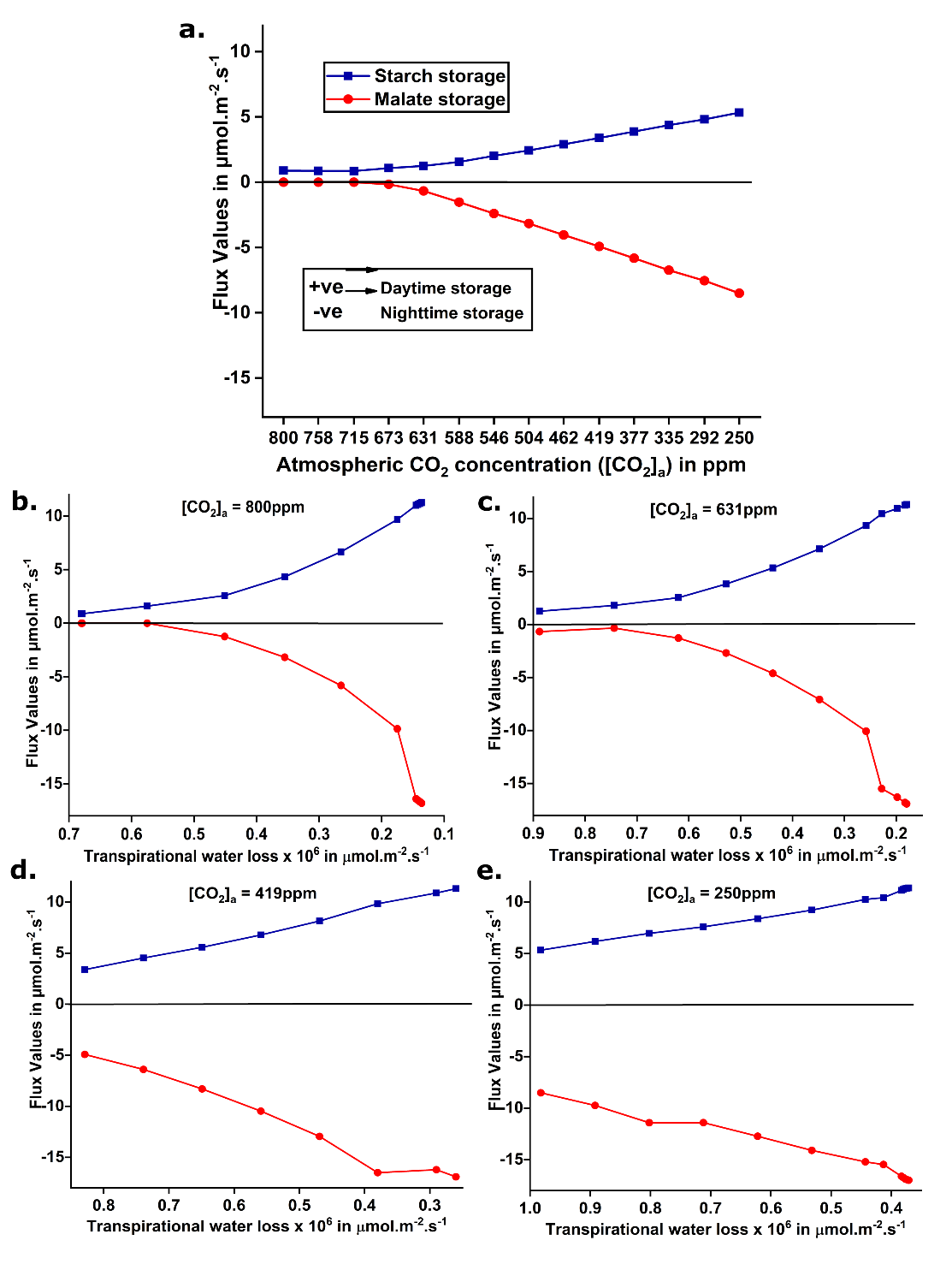

**Supporting Information Figure S5. Variation in starch-malate cycle. a** shows the increase in starch and malate storages at day and night respectively with the decline in [CO_2_]_a_; and **b-e** show the same with decreasing transpirational water loss at different [CO_2_]_a_. Only [CO_2_]_a_ values are shown in all the graphs. Corresponding T and RH values are listed in **Supporting Information Fig. S1c**. It should be noted that all the flux values presented in the graphs are normalized by phloem sap production.

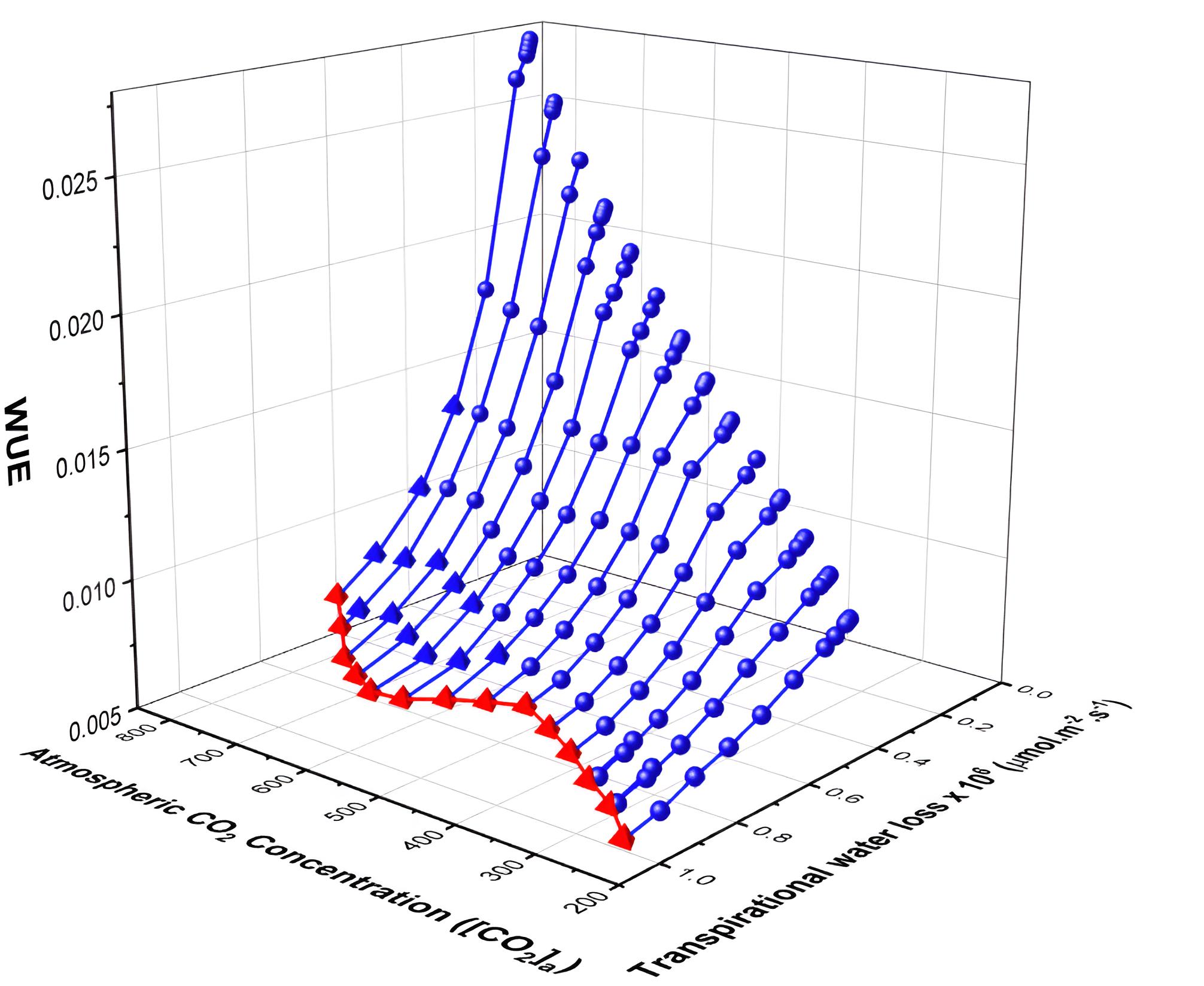

**Supporting Information Figure S6.** **The change in water-use efficiency (WUE) at different [CO_2_]_a_ and transpirational water loss.** The 3D plot shows the variation in WUE with [CO_2_]_a_ and transpirational water loss**.** Red colour represents the WUE values at different atmospheric conditions without any constraint on transpirational water loss. And the blue colour represents the WUE values at gradually reduced transpirational water loss in different atmospheric conditions. The tetrahedron and sphere indicate whether the model is representing C_3_ or CAM photosynthesis at a given point, respectively. We consider the metabolism as CAM, when there is a CO_2_ uptake at night. Only [CO_2_]_a_ values are shown in the graph. Corresponding T and RH values are listed in **Supporting Information Fig. S1c**.

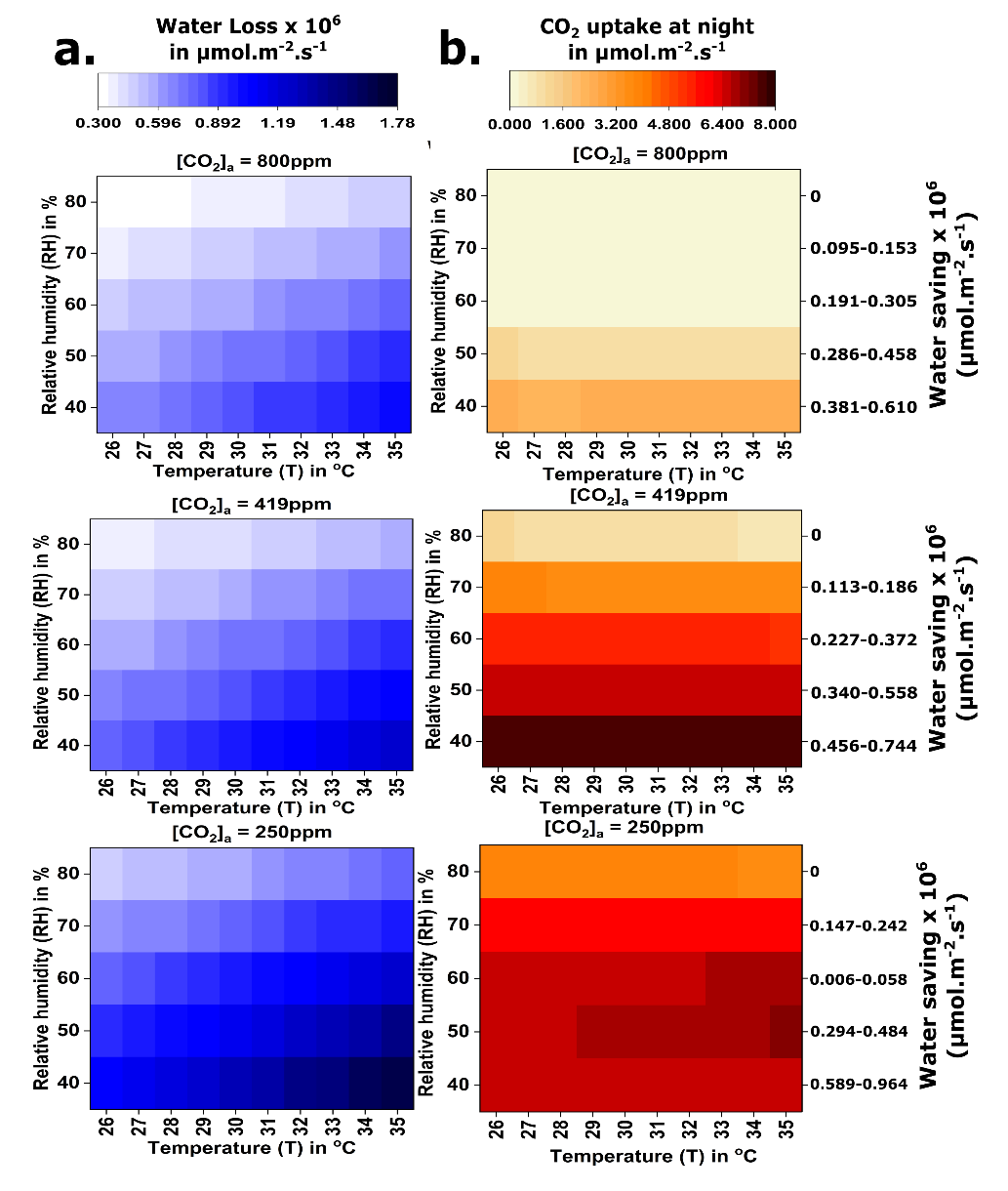

**Supporting Information Figure S7. Variation in transpirational water loss and nocturnal CO_2_ uptake at different T, RH and [CO_2_]_a_. a.** Variation in transpirational water loss in different combinations of [CO_2_]_a_, T and RH. Water loss increases with decrease in [CO_2_]_a_, RH and increase in T. **b** shows that for a fixed T and [CO_2_]_a_, maintaining constant transpirational water loss leads to increase in nocturnal CO_2_ uptake with decreasing RH, indicating the emergence of CAM under increasing aridity.

**Supporting Information Table S1.** Different constraints used for GC and MC throughout the C_3_-CAM transition.

| Cell Tpes | Constraints | Values |
| --- | --- | --- |
| GC | Ratio of light in the three phases of day | 5:148:3 |
|  | Nutrient uptake | 0 |
|  | Phloem sap production | 0 |
|  | Ratio of carboxylase and oxygenase activities of RuBisCO in each phase of day in C_3_ | 3:1 |
|  | Ratio of carboxylase and oxygenase activities of RuBisCO in each phase of day in CAM | 5.15:1 |
| MC | Ratio of light in the three phases of day | 5:148:3 |
|  | Ratio of nitrate uptake at day and night**^1^** | 3:2 |
|  | Rate of phloem sap production (in µmol.m^-2^.s^-1^) | 0.259 |
|  | Ratio of phloem sap production at day and night**^2^** | 3:1 |
|  | Percentage (%) of total accumulated osmolyte in GC to be sucrose, transported from MC to GC in phase 2 at daytime | 80 |
|  | Percentage (%) of total accumulated osmolyte in GC to be sucrose, transported from MC to GC in phase 5 at nighttime | 80 |
|  | Ratio of carboxylase and oxygenase activities of RuBisCO in each phase of day in C_3_ | 3:1 |
|  | Ratio of carboxylase and oxygenase activities of RuBisCO in each phase of day in CAM | 5.15:1 |

**^1^**The ratio is further distributed according to the time span of the phases

**^2^**It is free to be produced in any phase of the diel cycle

**Supporting Information Table S2.** Comparison of flux values through some specific reactions of CAM under different [CO_2_]_a_ values.

| **Properties** | **Flux values in CAM as observed in the last point of Fig. 1c at 250ppm**  **(µmol.m^-2^.s^-1^)** | **Flux values when CAM is simulated under elevated [CO_2_]_a_ of 500ppm**  **(µmol.m^-2^.s^-1^)** |
| --- | --- | --- |
| **Total daytime CO_2_ uptake** | 6.50 | 11.50 |
| **Total nighttime CO_2_ uptake** | 4.05 | 0.58 |
| **Daytime starch storage in MC** | 5.53 | 3.48 |
| **Nighttime malate storage in MC** | 8.82 | 4.90 |
| **Flux through PEPC at night in MC** | 8.54 | 4.15 |
| **Flux through MDH at night in MC** | 9.79 | 5.05 |
| **Flux through PEPCK at day in MC** | 7.34 | 3.84 |
| **Flux through ME at day in MC** | 1.54 | 1.54 |

**Increased flux of TCA cycle enzymes and O_2_ uptake with the emergence of CAM**

Results show that nocturnal mitochondrial respiration increases with the emergence of CAM. This is supported by elevated nocturnal O_2_ uptake, and increased flux values through several TCA cycle enzymes including 2-ketogluterate dehydrogenase (2KGDH), succinyl-CoA synthetase (SucCoASyn), succinate dehydrogenase (SDH), fumarase, and MDH (**Figure 2f**, **Supporting Information Dataset S4**). Among these, 2KGDH, SDH and MDH produce NADH and FADH_2_, which subsequently participate in mitochondrial electron transport chain (mETC) to produce higher amount of ATP. This increased production of mitochondrial ATP, essential for CAM cycle, also enhances the demand for the nocturnal O_2_ uptake. Additionally, simulation with restricted nighttime O_2_ uptake reveals that increased nocturnal O_2_ uptake is mandatory for CAM functioning, supporting the experimental observation that CAM requires elevated nocturnal respiratory rate and nocturnal O_2_ consumption (Leverett & Borland 2023). Interestingly, atmospheric O_2_ concentration increased during that geological time period (Berner, VandenBrooks & Ward 2007), which probably supports the higher O_2_ consumption and facilitated the emergence of CAM.

**Capturing possible changes in temporal differential activities of metabolic enzymes and transporters during early-Eocene to early-Miocene time period**

We calculate the Spearman’s rank correlation between the flux values through each of the enzymatic reactions and intracellular transporters across different cell types and the six phases of the diel cycle with- i) the gradual atmospheric changes during early-Eocene to the early-Miocene and ii) the gradually reduced transpirational water loss at each point between the same geological time interval (**Supporting Information Dataset S7**). The correlation values reveal a similar trend in the shift of the temporally differential flux distribution pattern in MC during the transition in both the cases. This pattern also aligns with our earlier findings during the C_3_ to CAM transition in reduced transpirational water loss at a fixed T, RH and [CO₂]ₐ (Sarkar & Kundu 2025). The enzymes showing gradual changes in MC are spanned across various metabolic pathways, including carboxylation-decarboxylation, glycolysis, gluconeogenesis, TCA cycle, pentose phosphate pathway, mitochondrial electron transport chain etc. However, GC exhibits different patterns, possibly due to the different stomatal behaviours. Under condition (i), as [CO_2_]_a_ declines, the plant’s CO_2_ demand necessitates wider stomatal opening (**Figure 3e, Supporting Information Figure S2**). As the stomatal pore has an upper physiological limit, very low [CO_2_]_a_ forces the plant to open their stomata both at day and night to meet the required carbon demand (**Figure 3e, Supporting Information Figure S2**). However, under condition (ii), at higher [CO_2_]_a_ (800 ppm), the minimization of transpirational water loss mainly drives the C_3_ to CAM transition, and a large number of reactions in GC show similar pattern as observed in our previous study (Sarkar & Kundu 2025). In contrast, at lower [CO_2_]_a_ (250 ppm), CO_2_ limitation alone significantly drives CAM-like features even without any restriction in water loss, and further reduction in water loss has minimal impact, resulting in lower water saving.

**Simulating the model considering all potential osmolytes and transfer metabolites**

In our model, we previously considered K^+^, Cl^–^, malate and sucrose as osmolytes to maintain the OP (Talbott & Zeiger 1996; Lawson 2009; Daloso et al. 2016; Daloso et al. 2017; Santelia & Lawson 2016; Robaina-Estévez et al. 2017) and allowed the transfer of only sucrose (Lawson, Simkin, Kelly & Granot 2014; Daloso *et al.* 2016) (80% of the osmolytes required for stomatal opening) from MC to GC in phase 2 and 5. Now, we have further included other potential osmolytes (nitrate, glucose, fructose and maltose) and transfer metabolites (glucose and malate) in our model (**Supporting Information Dataset S9**) based on other experimental studies (Talbott & Zeiger 1993; Lawson et al. 2014; Daloso et al. 2017; Flütsch et al. 2020b a; Dang et al. 2024). We have allowed the metabolites to transfer freely from MC to GC in any of the six phases of the diel cycle. Without any restriction on transport, GC takes its required carbon from MC and does not depend on its own photosynthesis. GC’s photosynthetic capacity is reported to be lower than MC (Lawson 2009); but it is not proved that GC doesn’t have any. Therefore, we constrain the transfer of glucose, malate, and sucrose from MC to GC in such a way that 80% of GC’s carbon demand is supplied from the MC. We imposed an additional constraint in phase 1, allowing GC to receive only up to 10% of the total carbon transported from MC. This adjustment helps us to incorporate comparatively lower activity of GC, as well as to observe the nighttime accumulation of starch in GC, which provides fuel for early-morning stomatal opening, in accordance with the experimental findings (Horrer *et al.* 2016). Without restricting metabolite transport in phase 1, GC would bypass the need for nocturnal starch storage. Now, with theses additional osmolytes and transfer metabolites, we simulate the model to check whether ancient atmospheric changes during early-Eocene to early-Miocene or restricted transpirational water loss at different points between that geological time period shift the C_3_ metabolism to CAM. Results again show that both aridity and declining [CO_2_]_a_ can drive C_3_-to-CAM transition (**Supporting Information Dataset S10)**. While, we observe variations in osmolytes accumulation in GC, MC shows similar metabolic patterns even in different scenarios. A key advantage of metabolic modelling is its ability to capture potential fluctuations in osmolyte accumulation that contribute to building and maintaining GC turgor pressure.

In C_3_, K^+^ accumulates in the early morning, accompanied by malate and Cl^-^ to balance the positive charge, consistent with previous experimental findings (Talbott & Zeiger 1996; Daloso *et al.* 2017; Lawson & Matthews 2020). Glucose and fructose also accumulate in phase 1, which matches with the experimental reports suggesting the accumulation of glucose for stomatal opening in *Arabidopsis thaliana* (Flütsch *et al.* 2020b) and their accumulation continues in phase 2. Maltose is seen to accumulate in phase 3 and the experimental findings suggest the accumulation of maltose in all the phases of day in *Vicia faba* when epidermal peels are exposed to blue light (Talbott & Zeiger 1993). Although sucrose is known to accumulate during later phases of the day in C_3_ plants to sustain OP (Talbott & Zeiger 1996), our model does not initially reproduce this trend. Interestingly, when maltose accumulation is blocked, sucrose accumulation emerged.

Whereas as we move towards CAM by declining [CO_2_]_a_, stomata start to open at night as well. Simulation of the model with [CO_2_]_a_ of 250 ppm, T of 22°C and RH of 30% (last point of the **Figure 1b**), shows that glucose and fructose storage increase from phase 5 to 6 and stored overnight, in agreement with experimental observation on *Kalanchoe fedtschenkoi* (Hurtado-Castano *et al.* 2023). K^+^ storage is observed in phase 6 (Lefoulon & Blatt 2024) and to balance the positive charge malate and Cl^-^ storages are observed. Moreover, when transpirational water loss is restricted, we observe C_3_-to-CAM transition at different [CO_2_]_a_. Nighttime storages of glucose, fructose and malate are observed in phase 5 and 6 (Hurtado-Castano *et al.* 2023). K^+^ storage increases at night (Lefoulon & Blatt 2024) and Cl^-^ and malate also accumulate to balance the positive charge of K^+^. Experimental studies also reported the accumulation of sucrose at night in CAM (Hurtado-Castano *et al.* 2023). Initially, we do not observe any night time sucrose storage; however, restrictions on glucose and fructose storage allow the guard cell to accumulate sucrose at night for maintaining the OP. These indicate that different combinations of osmolytes can maintain OP for stomatal opening under varying metabolic conditions, and such regulation likely involves not only metabolic but also other types of cellular control.

**Simulating C_3_ and CAM under predicted atmospheric changes in future**

One of the major reasons of CAM emergence is reduced [CO_2_]_a_. However, after the industrial revolution, [CO_2_]_a_ increased roughly by 40%, from a value of 278 ppm (Asadi *et al.,* 2017). It continues to increase day by day and is predicted to increase further in near future (Asadi et al. 2017). To understand the effect of this rising [CO_2_]_a_ on CAM, we simulate the model of CAM (as obtained in the last point of **Figure 2c**) under elevated [CO_2_]_a_. Results show that the higher [CO_2_]_a_ enhances the day time CO_2_ uptake, thereby reducing the reliance on the nocturnal CO_2_ fixation (**Supporting Information Table S2**), in agreement with the experimental findings reported by Sage *et al.* (2023). In future, not only [CO_2_]_a_, but temperature and aridity will also increase due to global climate change (Asadi et al. 2017) and it would be interesting to see how the plants respond in different combinations of [CO_2_]_a_, T and RH. Although we have already studied the dynamic temporal changes in plant metabolism with gradual change in atmospheric conditions from early-Eocene to early-Miocene (Fig. 1b), each simulation is done only under a particular combination of [CO_2_]_a_, T and RH. Therefore, we again simulate the model varying all the three atmospheric factors ([CO_2_]_a_: 800, 419 and 250 ppm, T: 26-35ºC with an increment of 1ºC and RH: 80-40% with a decrement of 10%) and observe that water loss increases with increase in T and decrease in [CO_2_]_a_ and RH (**Supporting Information Figure S7a**). For any fixed T and [CO_2_]_a_, transpirational water loss increases as RH decreases (or aridity increases) (**Supporting Information Figure S7a, Supporting Information Dataset S11**). But, in reality, increasing aridity limits the water availability (Wang et al. 2023) , and hence plants try to reduce the transpirational water loss for survival. At any T and [CO_2_]_a_, the transpirational water loss at 80% RH is the lowest (within the range of our simulation). To understand the effect of low water availability at increased aridity, for a fixed [CO_2_]_a_ and T, we simulate the model at reduced RH values, keeping the water loss fixed at a value obtained with RH of 80%. This constraint, representing water saving conditions, induces a shift towards CAM (**Supporting Information Figure S7b,** **Supporting Information Dataset S11**). Nighttime CO_2_ uptake and water saving increase with aridity when we impose a constraint on transpirational water loss. At higher [CO_2_]_a_ levels, water saving forces the plant to shift the time of CO_2_ uptake from day to night as temperature is comparatively lower at night. Here, larger water saving is possible without any compromise in phloem sap production. However, under low [CO_2_]_a_, even fully opened stomata throughout the day can’t capture enough CO_2_, causing the plant to take CO_2_ at night as well, even without any restriction in transpirational water loss (**Supporting Information Figure S7a**). Under this condition, a significant reduction in transpirational water loss (or water saving) reduces the phloem sap production to cope with the limited CO_2_ availability (**Supporting Information Figure S7a,** **Supporting Information Dataset S11**). This is in agreement with previous research which showed that photosynthetic productivity of C_3_ plants is reduced in low [CO_2_]_a_ (Sage et al. 2023), the productivity of CAM plant is lower than that of C_3_ (Shameer et al. 2018) and CAM plants have reduced phloem sap production compared to C_3_ due to the trade-off between water-saving and phloem sap production (Töpfer *et al.* 2020).
